# City climate and landscape structure shape pollinators, nectar and transported pollen along a gradient of urbanization

**DOI:** 10.1101/2021.10.15.464497

**Authors:** Paolo Biella, Nicola Tommasi, Lorenzo Guzzetti, Emiliano Pioltelli, Massimo Labra, Andrea Galimberti

**Affiliations:** University of Milano-Bicocca, Department of Biotechnology and Biosciences, ZooPlantLab, Piazza della Scienza 2, 20126, Milan, Italy

## Abstract

1. Urbanization gradients influence both landscape and climate and provide opportunity for understanding how species, especially plants and pollinators, respond to artificially driven environmental transitions.
2. Here, we investigated several aspects of pollination along an urbanization gradient in landscape and climate. We quantified wild hoverfly and bee abundances with trapping, standing crop of nectar with spectrophotometer, and the pollen transported by flower-visitors with DNA-metabarcoding, in 40 independent sites from seminatural to built-up areas in Northern Italy.
3. Linear and nonlinear relationships were detected along the urbanization gradient. Pollinator abundances increased until 22% of impervious surface, dropping by 34% after that, and it also decreased with green patch distance and urban park size. Thus, suburban landscapes host more pollinators than highly cemented or seminatural areas. Moreover, pollinators diminished by up to 45% in areas with low temperature seasonality: urbanized areas likely posing thermic stress. Furthermore, the sugar mass available in nectar increased by 91% with impervious cover, indicating that city nectars were less consumed or flowers more productive. Furthermore, the species richness of pollen decreased by 32% in highly urbanized areas, and with a high incidence of exotic plants, hinting for anthropized plant communities.
4. *Synthesis and applications*. Urbanization influences pollinator abundances, nectar resources and transported pollen. Pollinators are negatively affected by a thermally harsh climate in highly urbanized areas with isolated green areas and large parks. Suburban landscapes demonstrated the highest pollinator presence. In the city core, flowers contained more nectary sugar, while pollinators collected pollen from a small number of plants, mainly exotic. These findings highlight the influence of urban landscape structure and climate on pollinators and plants, showing that cities are heterogenous realities. Patterns from this study will serve as basis for pollinator-friendly planning and management of urban landscapes.

## Introduction

Given the environmental change caused by cemented and built-up surfaces in contrast to adjacent ecosystems, urban areas are often considered as a separate macrohabitat for animal and plant communities (Bolger *et al*. 2000; Faeth and Kane 1978). Cities provide an artificial environment that creates an extreme transition (Lemoine-Rodríguez *et al*. 2020), causing several types of responses on biodiversity. These impacts affect a number of ecosystem processes (McIntyre et al. 2001), mainly by influencing changes in species interactions (Cohen *et al*. 2020; Geslin *et al*. 2013). As plants and pollinators play key roles in many ecosystem processes (Patel *et al*. 2020; Potts *et al*. 2016), gaining an understanding of how urbanization gradients shape aspects of pollination and pollinator ecology is of utmost importance.

Urbanization can be described as a gradient, with different consequences for plants and pollinators. For instance, plant diversity is higher at moderate urbanization (McKinney 2008), and suburban areas host higher wild bee and butterfly diversity than the city core (Banaszak-Cibicka and Żmihorski 2020; Kurylo *et al*. 2020). Surprisingly, the potential beneficial aspects of suburban landscapes on pollinator and plant communities has rarely been pointed out to-date (Wenzel *et al*. 2020; Harrison and Winfree 2015). Moreover, different pollinator types and life-history traits respond differently to urbanization (Wenzel *et al*. 2020) in spite of a generally high variation among studies (Buchholz and Egerer 2020). For instance, less Diptera Syrphidae than Hymenoptera are expected to occur in cities (Persson *et al*. 2020). Within Hymenoptera, built environments may change the composition of bee assemblages, with a high occurrence of solitary and above-ground nesting bees (Wilson and Jamieson 2019), while filtering big species out (Buchholz and Egerer 2020). Furthermore, fragment isolation may play a role, as a study found poorer pollinator assemblages in more isolated urban green areas (Tonietto *et al*. 2011). All these aspects highlight the need to further explore how pollinators respond to structural differences in urbanized landscapes.

Cities transform not only landscapes but they also directly impact local climates (Kuttler 2008), thus likely triggering species physiological responses, even in plants and pollinators. Urban climate is are usually warmer, it holds lower relative humidity and has higher precipitation than the surroundings (Kuttler 2008). The urban heat island effect determines both high temperatures and lower temperature seasonality, a phenomenon that lasts both daily and across seasons (Marando *et al*. 2019). For instance, warmer cities impact plant physiology and phenology, triggering earlier flowering (Fisogni *et al*. 2020; Neil and Wu 2006). Also pollinator physiology can be affected (Hamblin *et al*. 2018). Intriguingly, a recent study showed that bees inhabiting city cores have higher thermic stress (Burdine and McCluney 2019). It is likely that the climatic impact posed by cities on plant and pollinator physiology are directly connected to patterns of diversity and abundances of those assemblages (Chown and Duffy 2015; Diamond *et al*. 2015). This is exemplified by studies showing that climatic features could affect some families of bees even more than landscape alteration (Kammerer *et al*. 2021). Therefore, it seems relevant to describe the responses of plants and pollinators not only as a function of city landscape alone but also by urban climatic variation.

In this study, we evaluated the effects of urbanization on several aspects of pollination and pollinator ecology, which are also relevant for ecosystem functioning (Biella, Akter, *et al*. 2019; Patel *et al*. 2020). We surveyed along a gradient of increasing urbanization in Northern Italy, a region characterized by a high proportion of built-up surfaces and remarkable climatic shift due to urban areas (Perini and Magliocco 2014). In order to address a mechanistic understanding of how an artificial gradient shapes pollination ecology and to connect local surveys to the structure of the surrounding area, we characterized landscape composition, its configuration and the climate.

We tested factors that were specifically selected based on hypothesized direct effects. Firstly, we measured pollinator abundance, as an indicator of habitat suitability (Bartholomée *et al*. 2020) and hypothesized that, along the urbanization gradient, pollinators may be dependent on the distribution and accessibility of suitable areas measured here as proportion of impervious cover and isolation of green spaces used for foraging (Steffan-Dewenter and Tscharntke 1999). Within the context of the highly urbanized area, we also tested the role of city park size in order to further highlight the relationship with patch size as larger areas might serve as refugia for larger populations (Baldock *et al*. 2019). In addition, temperature variation along the urbanization gradient could impact flower visiting organisms and also determine their local abundance (Burdine and McCluney 2019; Hamblin *et al*. 2018) by affecting pollinator physiology (Colinet *et al*. 2015). Secondly, we characterized the availability of nectar sugar mass, because nectar constitutes one of the main resources collected by flower visitors (Hicks *et al*. 2016). The reward quantity along the gradient would mainly be due to the secretion rate and thus to plant physiology, and its amount is partly determined by pollinator foraging rate (Corbet 2003). We hypothesize that the available nectar quantity could depend on the size of green areas because bigger patches may host richer communities(Collins *et al*. 2009; Dauber *et al*. 2010). In addition, plant productivity is often driven by precipitation and by the length of the thermally suitable season for growing, climatic parameters that vary with urbanization and that could impact nectar production (Mueller *et al*. 2020; Zipper *et al*. 2016). Thirdly, we considered the pollen diversity carried by flower-visitors, that were used here as a passive surveying of plant richness from the pollinator perspective (Biella, Tommasi, *et al*. 2019). In other words, we quantified the number of plant species visited by pollinators by studying the transported pollen, that is an important component of the total pollination rate (Bosch *et al*. 2009). Moreover, the advantage of using pollinators as passive samplers lies on the difficulty of sampling the vegetation of complex urban structures (e.g. tall buildings, balconies and private gardens) that impede traditional survey techniques. Here we described the transported pollen richness along the urbanization gradient (hence in relation to the impervious cover) and we also expected an influence by green-patch sizes, as it may determine local plant diversity (Collins *et al*. 2009; Dauber *et al*. 2010). We did not analyze climatic variables in this case as in anthropogenic areas pollen availability could be influenced by management (Aronson *et al*. 2017; Ibsen *et al*. 2020).

## Materials and methods

### Study area

Study sites were set in Northern Italy, mainly in the region surrounding Milan (Fig. 1) that is occupied by urbanized surfaces (about 38% of the area), intensive agricultural environments (ca. 53%), and natural forests, wet habitats and seminatural hay meadows (ca. 9%) (Regione Lombardia and ERSAF 2010). Study site locations were selected randomly with a GIS software (QGIS 3.6.2) and were distributed in an area of 1575 km^2^ over a surface including the entire region of Milan and the urban parks of Milan city. We applied a minimum of 1 km distance between points to assure the independency of sampling sites (Phillips *et al*. 2019), later confirmed by an autocorrelation analysis resulting not significant when testing correlation of putative predictors with themselves with Moran”s I (P value > 0.05). We adjusted the exact sampling locations so that each would be located in either a urban park surrounded by “impervious” surfaces (*i.e*., concrete, asphalt, buildings) or at the margins of agricultural fields with a varying quantity of surrounding impervious surface or in seminatural hay meadows near forests (<1 km) with little amount of urbanization nearby (Fig. S1). Overall, 40 sites were surveyed, across the entire urbanization gradient (Fig. 1, Fig. S1).

**Figure 1.**
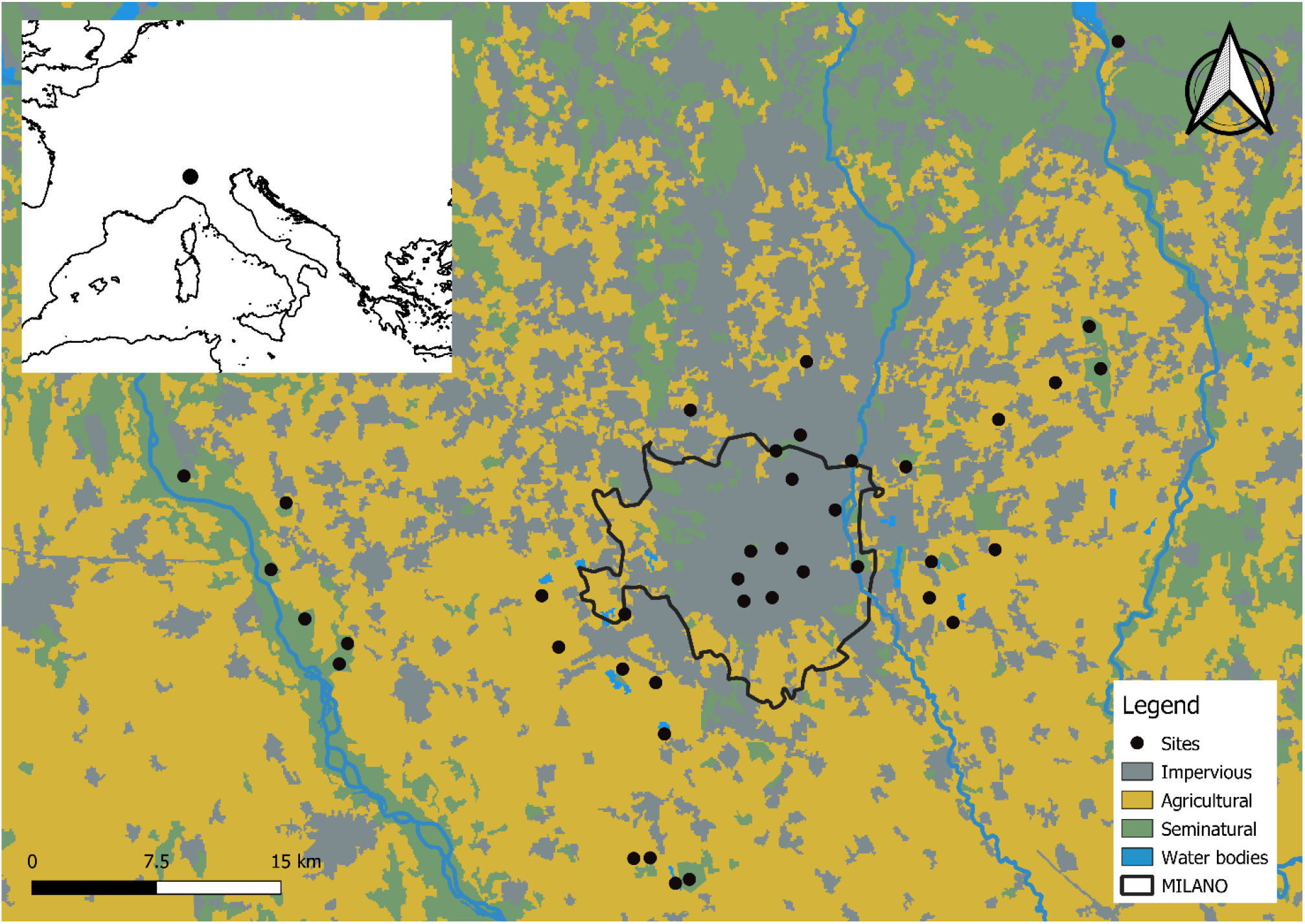
Sampling sites. The distribution of sampling sites along a gradient of increasing urbanization in Northern Italy. The map also shows the Municipality of Milan, three major rivers and the main land uses of this region.

### Landscape and local climate characterization

To characterize landscape composition, we measured the cover percentage of impervious (concrete-dominated) surfaces, urban green areas and seminatural lands (forests and hay meadows). To describe landscape configuration we focused on green and seminatural patches together and calculated Edge density (ED), the Euclidean nearest neighbor distance (ENN), and also the size of urban parks. ED measures the total length of all edge segments per area unit and describes the fragmentation in size of the patches (*i.e*., highly fragmented when the perimeter is high relative to their area), while ENN describes the mean distance between patches in the landscape.

Temperature and precipitation vary with urbanization (see Figs S2-S3), we used 3-months averages of hourly temperature at 2-m and hourly precipitation sum between February and July 2019. We calculated an index of temperature seasonality as the coefficient of variation between spring and summer. Air temperature data were also used to calculate the Growing Season Suitability index (GSS), quantifying the length of the vegetative season that, together with precipitation, could impact the productivity of plant resources. All used variables were extracted from circular buffers of 1 km around the sampling sites, further methodological details are available in Appendix A1.

### Pollinator abundance

Sampling of pollinator abundances took place during the peak flowering period for the study area, from mid-May to mid-June 2019, in 36 of the original 40 sites. Passive sampling took place for 24 hours at each site, with pan and sticky traps, that are complementary methods for sampling the same target insect groups. The pan trap consisted of a wooden stick holding three pan traps of yellow, blue and white bowls with UV reflection, containing water with 1-2 mL of soap to reduce surface tension, and the bowls height was adjusted specifically depending on grass tallness, so that the trap would be slightly higher than the height of the grass (Popic *et al*. 2013). Sticky traps were 15 * 20 cm yellow plastic board placed on poles at 1 m height on which pollinating insects remain glued (Sutherland *et al*. 2001). In each site, three sets of pan-traps were placed 10 m apart from each other; and five yellow sticky traps were placed at 5 m from the pan-traps. In this study, we counted the total abundance of bees and hoverflies (Hymenoptera: Anthophila and Diptera: Syrphidae) following (Bates *et al*. 2011). We did not count honeybees since their numerosity is only due to beehives in the vicinities. In the statistical analyses that followed we kept wild bees and hoverflies together and evaluated their total abundance as they belong to the same guild of pollinators (see Fig. S4).

### Nectar sugar quantification

Standing crop of sugar mass in the nectar was measured from mid-May to mid-June in 35 sites. Flowers of two to three most abundant herbaceous flowering species were selected after an inspection of plant relative cover based on the number of stalks. We chose abundant plants as they should offer most of the local resources. Standing crop of nectar sugar mass is a measure of nectar quality available at a given time and it is a function of both plant secretion and of pollinator visitation frequency (Corbet 2003). As in Biella, Akter, et al. (2019), twenty to thirty blooming flowers for each species were taken from individual plants and the internal part of the corolla was washed in distilled water with a 100 μl Hamilton syringe, the number of processed flowers was noted, samples were weighed and then frozen. Sugar quantification was performed using a spectrophotometer Cary 60 (Agilent Technologies, USA) and with Sucrose, D-Fructose and D-Glucose kits (Megazyme, Ireland). We used the sugar mass per flower for each given species in subsequent statistical analyses, which was calculated by dividing the sugar mass by the number of washed flowers of a species processed at a site (Biella, Akter, *et al*. 2019).

### Pollen richness with DNA metabarcoding

In each sampled site, insects foraging on flowers were actively sampled for one hour by hand-netting and then stored in sterile Eppendorf tubes filled with ethanol 70%. In the laboratory, we randomly chosen a subset of the sampled sites (N=25) and analyzed the pollen from insect bodies with DNA metabarcoding following the protocol of Biella, Tommasi, et al. (2019). Full details of the laboratory protocol and bioinformatic processing are reported in Appendix A2. For each site, the number of plant species found in the pollen of all flower visitors was used as indication of pollen richness.

### Statistical analyses

Putative predictors were chosen based on the ecological hypotheses outlined above. We used a repeated K-fold cross-validation for choosing if to fit linear or non-linear models by selecting the lowest RMSE (Root Mean Squared Error) between a linear GLM (Generalized Linear Model) and a non-linear GAM (Generalized Additive Model) associated to each predictor, ten repetitions of K=10 were performed and the mean RMSE was used for the evaluation (see Table S1). To exclude collinear variables from the same analyses, we calculated the VIF index from preliminary regression models (Variance Inflation Factor, with an exclusion threshold of 4, Table S2) with the R package *car* (Fox and Weisber 2019). In the regression models listed below, predictors were square rooted to correct variable skewness and they were scaled to avoid different numerical ranges. Statistical significance was tested with likelihood-ratio tests. We analyzed the effects of landscape composition and configuration, and the summer-to-spring temperature seasonality on pollinator abundances in a Generalized Additive Model with the proportion of impervious land fitted with a smooth term, and the ENN and temperature variation as linear terms. These predictors were chosen because impervious cover describes the urbanization, the ENN indicates distance between patches used by pollinators, and temperature seasonality indicates the potential for thermic stress. This regression was fitted with Structural Equation Models (SEM), including the correlated errors between all predictors (Table S2), with the *piecewiseSEM* package with R (Lefcheck 2016). In addition, we modelled the relationship between pollinator abundances and urban park size with a GLM with data from urban parks (this predictor was log-transformed). In all cases, a Poisson family was used for error distribution with a log link function.

For sugars mass, we analyzed the effects of landscape variables and the contribution of urban climate. In particular, we fitted the sugar mass per flower as response and the proportion of impervious land, ED, the mean summer precipitation and the GSS as predictors in a Generalized Linear Mixed Model (GLMM), with a Gamma error distribution, the logarithm as a link function, and plant species and site identities as random intercepts. While impervious cover characterize the urbanization, the other variables describe the impact on plant productivity as ED explains the role of patch size on the diversity of plant communities, and the mean summer precipitation and GSS measure the amount of natural watering and length of the favorable season. This regression was fitted with Structural Equation Models (SEM), including the correlated errors between all predictors (Table S2).

We also analyzed the effect of the urban landscape on the transported pollen richness. We used the total pollen richness for each site as response variable in a GLM with the proportion of impervious land and ED as predictors, a Poisson family for error distribution and a log link function. These variables were used because impervious cover indicates the urbanization gradient and ED considers the role of patch size in hosting plant diversity, while we did not expect a contribution from seasonal climate on the transported pollen. In addition, we investigated the connection between the geographical origin of plants and the landscape on the plant species of pollen transported. To do so, we used the Fourth Corner analysis to evaluate the relationship between plant incidence measured as the proportion of samples where a plant was found at a given site, plant traits as native of the sampled region, agricultural crop or exotic (Galasso *et al*. 2018), and site attributes as the proportion of impervious, seminatural and urban green cover. The analysis was performed with *ade4* package in R (Thioulouse *et al*. 2018) by setting 999 permutations of sites and species values for testing significances.

## Results

Pollinator abundances were nonlinearly dependent on the landscape variable of impervious cover, while it decreased linearly with green patches ENN (Table 1, Fig. 2). Pollinator abundance was positively linked to impervious cover until a threshold value of 22% cover after which the relationship became negative, decreasing by 34% (Fig. 2A). Increasing green patches ENN were associated with a decline in pollinator abundance of 27% across the range of the variable (Fig. 2B). Abundances of pollinators positively responded in a linear way to the seasonal variation in mean temperature between spring and summer (Fig. 2C), with the highest thermic seasonality in sites with medium to low impervious cover (see Fig. S2). Specifically, pollinators increased by 45% across the range of temperature variation. In city parks, the relationship between pollinator abundances and park size was negative (Table 1, Fig. S5), with a decline of 60% over the range of the studied urban parks (0.02 to 0.68 km^2^).

**Figure 2.**
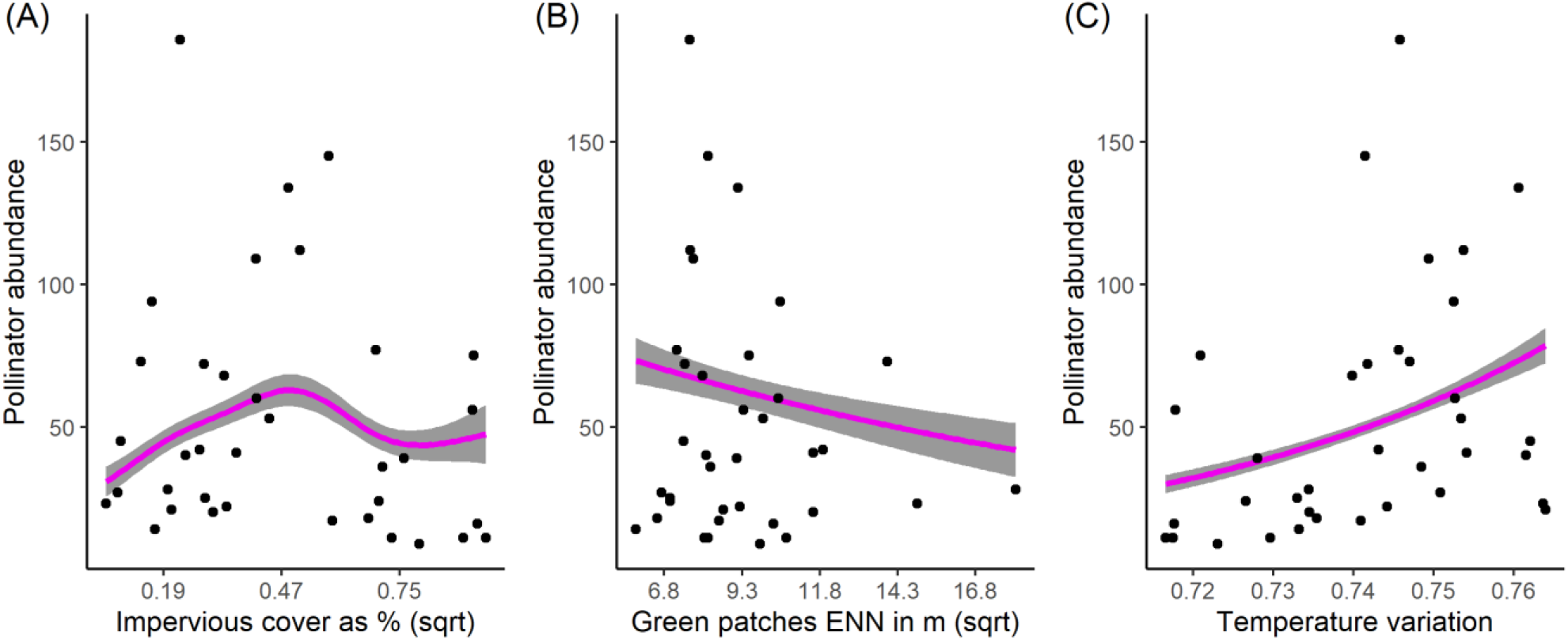
Pollinator abundances along an urbanization gradient. Relationship of pollinator abundance with (A) impervious cover as a percentage of buffer area and (B) the “ENN”, mean distances of green patches; these predictors were square rooted (“sqrt”). (C) Relationship of pollinator abundances with temperature seasonality between spring and summer. Statistical outputs are reported in Table 1.

**Table 1.** Estimated relationship between pollinator abundances, nectar sugar mass available and pollen species richness carried by pollinators and landscape and climatic variables. Significances are obtained with likelihood-ratio tests. SEM stands for Structural Equation Model; ENN indicates the Euclidean Nearest Neighbor distance of the green and seminatural areas; ED is the Edge Density, total length of all edge segments per unit of area; Growing Season Suitability index is indicated with GSS and it is the fraction of days between February and April with mean temperature above 10°C.

| Response | Model type | Predictor | Regression slope ($B_i$) | Degree of freedoms; Chi squared $\chi^2$ | Significance P |
| --- | --- | --- | --- | --- | --- |
| Pollinator Abundances | Urbanization landscape and climate model (SEM) | Impervious land | Smoothed, see Fig. 1 | 3.67; 57.30 | **< 0.001** |
|  |  | ENN | - 0.11 | 1; 16.25 | **< 0.001** |
|  |  | Temperature seasonality | 0.29 | 1; 58.36 | **< 0.001** |
| Pollinator Abundances | Urban parks (GLM) | Recreational park size | - 0.532 | 1; 81.467 | **< 0.001** |
| Nectar sugar mass | Urbanization landscape and climate model (SEM) | Impervious land | 0.89 | 1; 6.72 | **< 0.01** |
|  |  | ED | -0.37 | 1; 2.29 | 0.13 |
|  |  | Summer precipitation | 0.18 | 1; 0.92 | 0.33 |
|  |  | GSS | -0.25 | 1; 1.62 | 0.20 |
| Pollen richness | Urbanization landscape (GLM) | Impervious land | -0.23 | 1; 11.29 | **< 0.001** |

Sugar mass per flower was linearly dependent on impervious cover and on precipitations (Table 1, Figs 3A-B). Specifically, sugar mass increased by 91% across the range of impervious cover. However, it was not significantly dependent on green areas ED, or GSS or precipitation.

**Figure 3.**
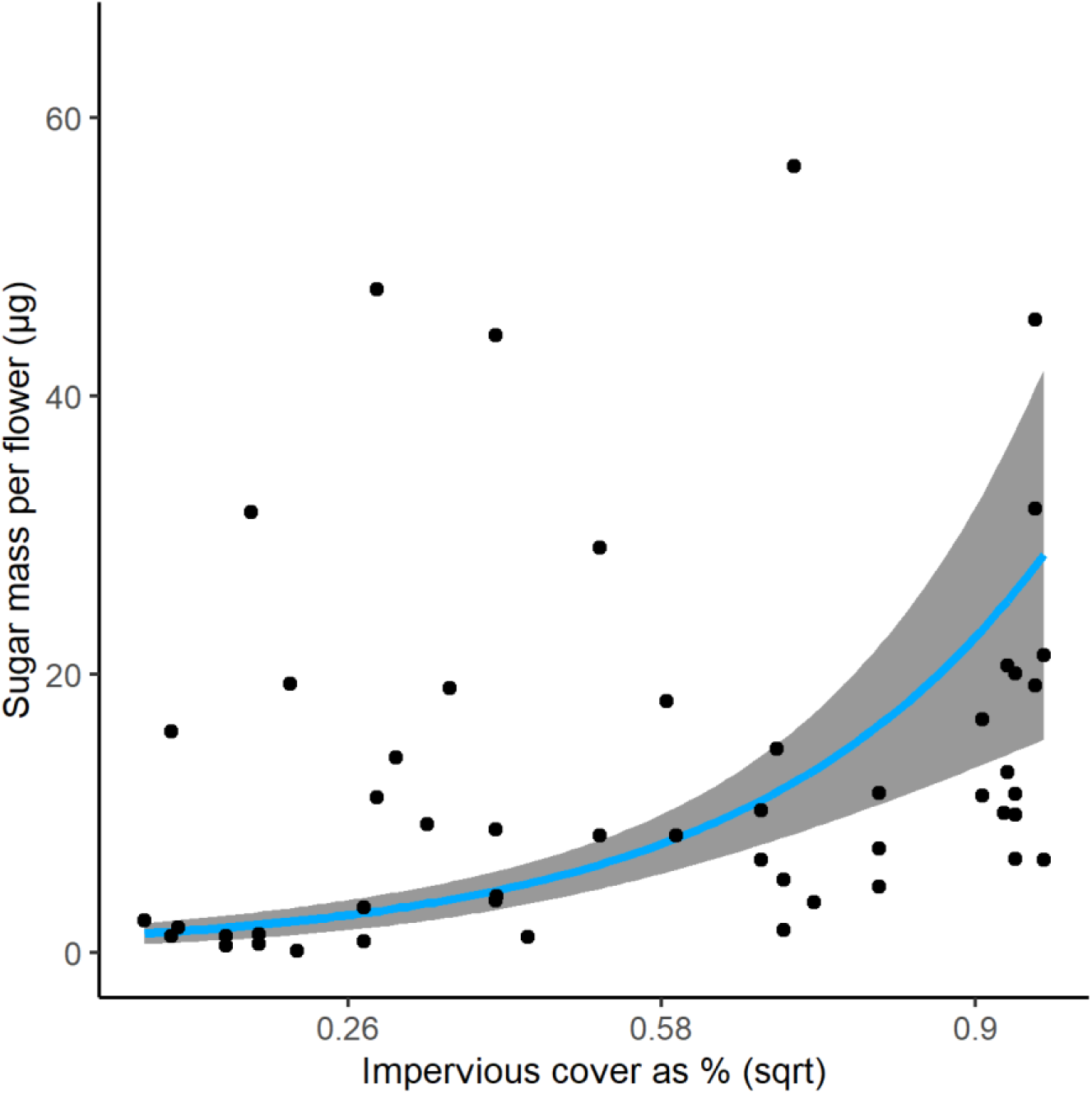
Sugar mass in nectars along an urbanization gradient. The relationship of sugar mass (standing crop) per flower with impervious cover as a percentage of buffer area is shown. This predictor was square rooted (“sqrt”). The statistical outputs are in Table 1.

The transported pollen richness showed a linear negative relationship with the impervious cover, and specifically by 32.5% across the range of the variable, and it was not significantly dependent on green areas ED (Table 1, Fig. 4). Most of the species in the pollen transported by pollinators were native to the region (66.1%). The incidence of exotic plants in the pollen samples was significantly higher at sites with higher cover of urban green areas (Table 2).

**Figure 4.**
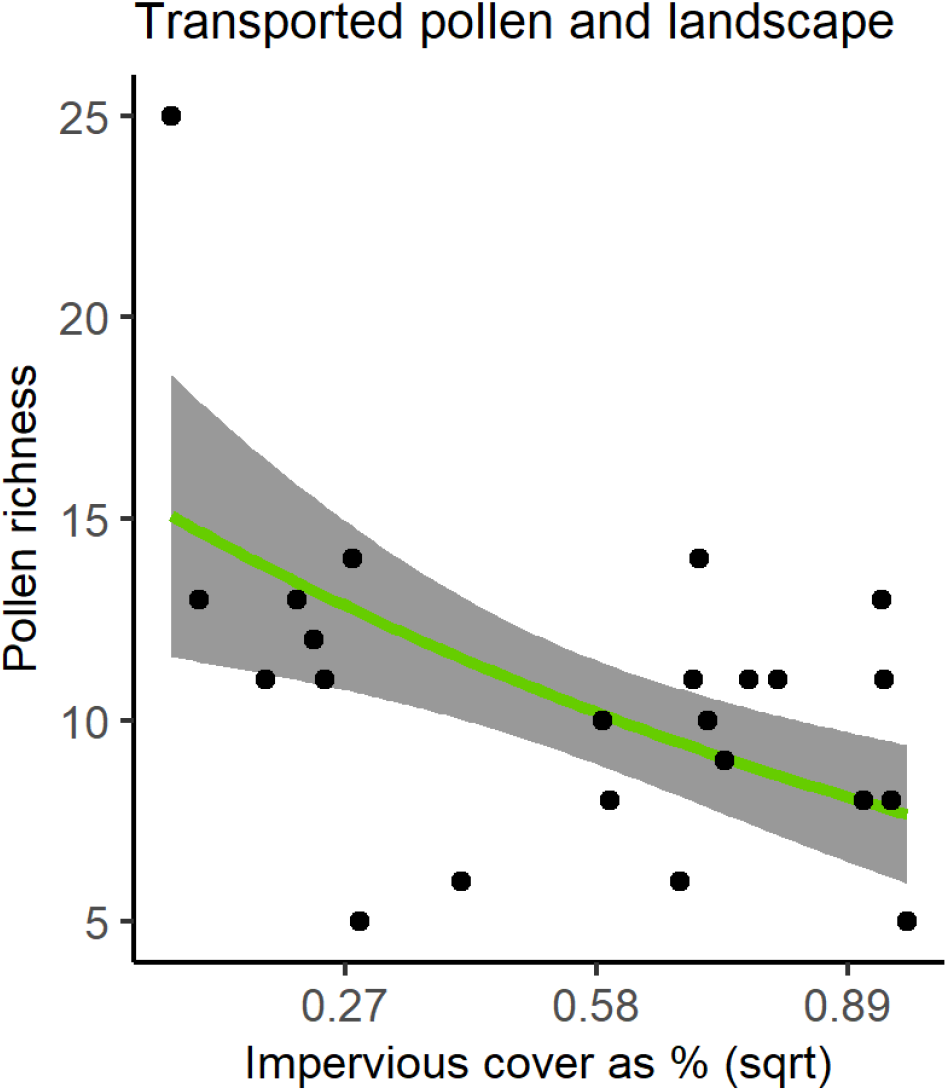
Plant species richness in pollen from flower-visitor along an urbanization. The relationship between pollen richness for each site and impervious cover measured as surface percentage, in circular buffers of 1 km. This variable was square rooted (“sqrt”). The statistical outputs are indicated in Table 1.

**Table2.** The relationships between the type of plant species found in the pollen and landscape composition. Plants are indicated as “Native”, “Crops” or “Exotic”. Results of a Fourth corner analysis are shown, based on the frequency of plant presence in pollinator samples of each site.

| Land use cover | Plant origin | Fourth corner statistic r | P value |
| --- | --- | --- | --- |
| Green areas | Exotic | 0.203 | **0.025** |
| Impervious | Exotic | 0.139 | 0.205 |
| Seminatural | Exotic | -0.194 | <u>0.077</u> |
| Green areas | Native | -0.156 | 0.115 |
| Impervious | Native | -0.076 | 0.499 |
| Seminatural | Native | 0.188 | <u>0.091</u> |
| Green areas | Crop | -0.036 | 0.696 |
| Impervious | Crop | -0.074 | 0.432 |
| Seminatural | Crop | -0.025 | 0.77 |

## Discussion

Along the gradient of urbanization, landscape composition impacted pollinator abundance in non-linear ways. At low-to-medium levels of impervious cover, the pollinators were increasingly more abundant with urbanization, but landscapes with impervious surfaces higher than 22% in cover negatively impacted pollinator abundance. This threshold is comparable with another study from North America where butterfly abundance decreased when more than 25% of impervious cover occurred (Kurylo *et al*. 2020). Thus, cities are not homogenous entities, but some parts of the urban gradient may benefit pollinators (i.e. suburban areas) while others such as the heavily urbanized city cores do not, a pattern already observed in other studies (Banaszak-Cibicka and Żmihorski 2020; Buchholz and Egerer 2020). Interestingly, pollinators were lowly abundant even at very low impervious cover, perhaps either due to pollinator-unfriendly practices (e.g. agrochemicals, mechanical disturbance, low wildflower cover and diversity) or because pollinator individuals are diluted over large open areas (Holzschuh *et al*. 2016). Moreover, the isolation of green patches negatively influenced pollinator abundances, in a linear way. This clearly indicates that the dispersion of green patches across a landscape may severely impact local pollinator abundances. This result recalls other studies showing that small and medium sized bees forage at close vicinities, for instance at maximal distance of 150 m (Hofmann *et al*. 2020; Zurbuchen *et al*. 2010). However, these studies were conducted in open landscapes, but in a urban setting it may be reasonable to expect lower home ranges given the presence of vertical obstacles (Harrison and Winfree 2015). Not only the type of landscape but also the size of city parks impacted pollinator abundances, as we recorded fewer pollinators with increasing park size. This result is comparable to what previously found in UK and in Germany (Baldock *et al*. 2019; Daniels *et al*. 2020). This may be due to low habitat quality (Bates *et al*. 2011) or a low population size diluted over a large area (Holzschuh *et al*. 2016). In spite of all these relationships, local conditions may play an important role in mitigating negative landscape impacts when nesting possibility and flowering resources are high (Delaney *et al*. 2020; Tommasi *et al*. 2021).

Temperatures also varied along the artificial gradient. Our analyses showed that pollinator abundances increased linearly with a rising temperature seasonality between spring and summer. This result indicates that pollinators are less abundant in sites where the climate is less variable between those seasons, and it contradicts previous ideas suggesting a link between a stable urban climate and pollinators (Baldock 2020). This is corroborated by a previous physiological study showing that wild bee species are affected by a high temperature where the impervious cover is high (Burdine and McCluney 2019). Furthermore, another study showed that bees avoid warmer areas in cities (Hamblin *et al*. 2018) and that warming may also reduce foraging activities (Kühsel and Blüthgen 2015). All together, these studies and our research indicate that pollinators may be sensitive to the harsh urban climate.

The impervious cover was positively associated with the standing crop of sugar mass available in the nectar of locally abundant plants, although independently to the fragmentation of the green areas (ED). These results showed a higher sugar mass available in urbanized areas than in non-urban sites. This result could be due to either higher secretion rate by plants or due to lower consumption by pollinators in cities (Corbet 2003). The latter possibility seems reasonable given a lower pollinator abundance in the city core that may translate into a lower consumption rate. Conversely, if future actions will increase pollinator abundances even in the core of the city, the foragers will consume more nectar, and this will likely modify the observed trends with impervious surfaces. As it seems that a higher proportion of sugar mass available in cities is a prominent feature of a highly urbanized landscape hosting few pollinators, conservation plans aiming to increase pollinator abundance or richness would impact the nectar pattern observed. It may follow that, as cities provide higher sugar mass, they would sustain high pollinator abundances if the urban landscape was more pollinator friendly. Another relevant aspect to highlight is that nectar availability could vary with plant phenology. Thus, the nectar pattern we observed might be altered by different plants being in flower due to seasonal phenology (Hicks *et al*. 2016). Plant phenology might even cause seasonal gaps of nectar resources with severe implications for pollinators (Timberlake *et al*. 2019), and thus our results should be interpreted within the time-frame of our investigation.

In addition to the described patterns in pollinator abundance and nectar, a negative and linear relationship was detected between the impervious cover and the richness of plant species found in the pollen collected by pollinators. This means that the pollen from fewer plant species were transported by the pollinators during their foraging trips in highly urbanized sites. As pollen diversity on flower visitor bodies often reflects the local flowering plant diversity (Biella, Tommasi, *et al*. 2019; Bosch *et al*. 2009), this result reveals that urban parks of the study area are currently not offering to pollinators as diverse plant resources as areas outside the city. This result could be connected to the low plant diversity usually found in highly urbanized areas (McKinney 2008; Wittig and Becker 2010). Interestingly, we detected a higher incidence of non-native pollen in sites with a higher cover of urban green areas. This indicates that the urbanization deeply shaped the foraging patterns of pollinators, which more frequently visited exotic flowers. Concerningly, pollinators of urban areas carrying less diverse pollen richness, dominated by non-native species, may even have direct implications for plant reproduction (Cohen *et al*. 2020).

In this study, the urbanization gradient set important scenarios for understanding how plant and pollinators respond to habitat alteration and environmental transitions of urban landscape and climatic features. We detected that the artificial gradient shaped pollinator abundances, pollen species richness transported by flower-visitors and sugar mass available in nectar in linear and nonlinear ways. These factors could have effects on plant reproduction, and on pollinator survival and nutrition. Importantly_this study clarifies that suburban areas, generally characterized by cemented surfaces at medium-low density and green patches of low isolation, host a high pollinator abundance. It is important to note that pollinator abundances are often correlated to species richness (Vereecken *et al*. 2021), and thus it could be expected to find similar patterns when considering also pollinator richness. However, highly urbanized areas provide nectars richer in available sugars, while the pollen transported was less rich of plant species and frequently with non-native plants, compared to less urbanized areas. As the gradient is human-driven, future actions could modify the responses observed in this study. In particular, managing green areas incorporating practices that are more pollinator-friendly will likely increase pollinator abundances and their activity (Turo and Gardiner 2019). Thus, increasing the suitability of existing and future urban landscapes for plants and pollinators is a priority, given the relevance played for ecosystem services and even for human health (Smith *et al*. 2015).

## Acknowledgements

The authors thank local authorities for the sampling permits. The authors also thank Giulia Agostinetto, Davide Magnani, Carola Miuccio, Aidana Nurtaza, Isabel Rondi and Andrea Tapparo for their technical help, and Kathryn Harrold for linguistic revision. This survey was funded by the PIGNOLETTO project, co-financed with the resources of POR FESR 2014-2020, European regional development fund with the contribution of resources from the European Union, Italy and the Lombardy Region. The funder had no role in conducting the research and/or during the preparation of the article.

## Author contribution

PB, ML and AG conceived the ideas and designed methodology; PB, NT, EP, LG collected the data; PB analyzed the data; PB led the writing of the manuscript. All authors contributed critically to the drafts and gave final approval for publication.

